# Detecting episodic evolution through Bayesian inference of molecular clock models

**DOI:** 10.1101/2023.06.17.545443

**Authors:** John H Tay, Guy Baele, Sebastian Duchene

## Abstract

Molecular evolutionary rate variation is a key aspect of the evolution of many organisms that can be modelled using molecular clock models. For example, fixed local clocks revealed the role of episodic evolution in the emergence of SARS-CoV-2 variants of concern. Like all statistical models, however, the reliability of such inferences is contingent on an assessment of statistical evidence. We present a novel Bayesian phylogenetic approach for detecting episodic evolution. It consists of computing Bayes factors, as the ratio of posterior and prior odds of evolutionary rate increases, effectively quantifying support for the effect size. We conducted an extensive simulation study to illustrate the power of this method and benchmarked it to formal model comparison of a range of molecular clock models using (log) marginal likelihood estimation, and to inference under a random local clock model. Quantifying support for the effect size has higher sensitivity than formal model testing and is straight-forward to compute, because it only needs samples from the posterior and prior distribution. However, formal model testing has the advantage of accommodating a wide range molecular clock models. We also assessed the ability of an automated approach, known as the random local clock, where branches under episodic evolution may be detected without their *a priori* definition. In an empirical analysis of a data set of SARS-CoV-2 genomes, we find ‘very strong’ evidence for episodic evolution. Our results provide guidelines and practical methods for Bayesian detection of episodic evolution, as well as avenues for further research into this phenomenon.

## 1. Introduction

Episodic evolution occurs when evolutionary change is accelerated over brief periods of time, such as through short-lived mutational bursts (Gillespie, 1984). Many microbial pathogens are known to undergo such periods of increased molecular evolutionary rates. Notably, the large number of mutations in SARS-CoV-2 variants of concern (VOC) is explained by these dynamics (Hill et al., 2022, Lythgoe et al., 2022, Markov et al., 2023, Neher, 2022, Tay et al., 2022). Similar processes are known to occur in other viruses, including during withinhost evolution in influenza viruses (Xue et al., 2018), in HIV lineages subject to hypermutation mediated by APOBEC3G/G (Simon et al., 2005), and upon host species jumps in SARS-CoV-2 (Porter et al., 2023). A major consequence of evolutionary rate acceleration is that it can result in increased transmissibility and virulence, as appears to be the case in some Ebola virus (EBOV) lineages (Mbala-Kingebeni et al., 2019) and SARS-CoV-2 VOCs, or drug resistance, which has been suggested for *Mycobacterium tuberculosis* (Cohen et al., 2015).

An effective method to detect episodic evolution using genome data is to estimate evolutionary rates for branches in phylogenetic trees using molecular clock models (Rannala and Yang, 2007, Yang and Nielsen, 2000). Here, the ‘evolutionary rate’ refers to the number of substitutions per site per unit time, such that it represents the combination of the instantaneous mutational process and the rate at which mutations are fixed (i.e. substitutions). Under this framework, branches that correspond to lineages expected to have undergone episodic evolution should therefore exhibit higher evolutionary rates.

The simplest molecular clock model is the strict clock (SC), where all branches are assigned the same evolutionary rate. The most flexible approaches, involve an uncorrelated ‘relaxed molecular clock’, in which evolutionary rates can vary across branches in the tree according to an underlying statistical distribution. For example, evolutionary rates across branches can be independent draws from a lognormal or gamma (Γ) distribution (typically abbreviated as UCLD and UCGD, for an uncorrelated clock with a lognormal and gamma distribution, respectively; reviewed by Ho and Duchene 2014 and Guindon 2020). An explicit model for episodic rates is the fixed local clock (FLC) (Worobey et al., 2014, Yoder and Yang, 2000). In the FLC model a set of branches, known as the *foreground*, are allowed to have a different evolutionary rate to the *background* (Fig. 1). To describe episodic evolution in the emergence of SARS-CoV-2 VOCs, the stem branches (i.e. between the stem and crown node) leading up to VOC clades are designated as the foreground (Gräf et al., 2021, Tay et al., 2022) (see also Fig. 1C).

**Figure 1:**
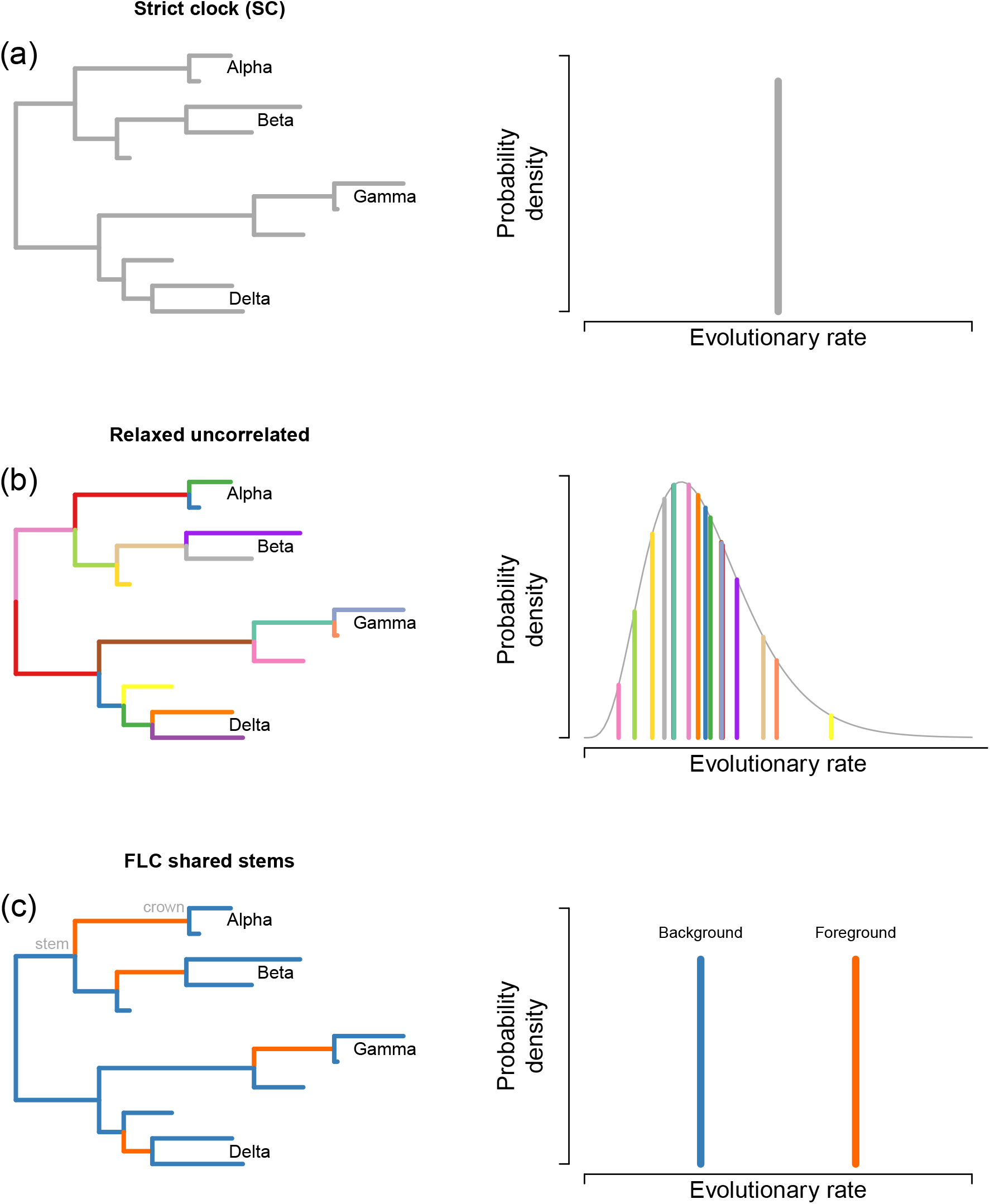
Examples of molecular clock models. (a) is a strict clock model (SC), where all branches share a single evolutionary rate. (b) is an uncorrelated relaxed clock model, in which the evolutionary rates across branches are independent draws from an underlying statistical distribution, such as a lognormal or a gamma (Γ) distribution (typically abbreviated as UCLD and UCGD, respectively). Note that branches are coloured according to those drawn from the distribution on the right. (c) represents a fixed local clock model, where stem branches leading up to SARS-CoV-2 variants of concern (VOCs; those leading to clades labelled as Alpha, Beta, Gamma and Delta, following variant names) are designated as the ‘foreground’ and assigned a rate that differs from the ‘background’. In (c), we have labelled the stem and crown nodes of variant Alpha, where evolutionary rate changes would have occurred.

Bayesian methods for quantifying the strength of evidence for episodic evolution include formal model testing. For example, one can compare the fit of the FLC with that of the SC and UCGD by comparing their associated (log) marginal likelihoods, allowing one to compute (log) Bayes factors of one clock model over the others, referred to here as ‘log Bayes factors on model comparisons’. Molecular clock models are treated as explicit hypotheses, with the UCGD and SC presenting a form of ‘null’ and the FLC the hypothesis of interest. Here, the log Bayes factor is the difference in log marginal likelihoods (i.e. the average fit) between competing models (Sinsheimer et al., 1996). There are multiple methods for estimating log marginal likelihoods in Bayesian phylogenetics (Lartillot and Philippe, 2006, Oaks et al., 2019), with generalised stepping-stone sampling being the most accurate to date (Baele et al., 2016, Fourment et al., 2020).

Following Kass and Raftery (1995), a log Bayes factor of two competing models, *M*_1_ and *M*_2_, 0 *< logBF*_12_ ≤ 1 is considered as not worth more than ‘bare mention’ for evidence of *M*_1_ over *M*_2_. A 1 *< logBF*_12_ ≤ 3 is considered as ‘positive evidence’, 3 *< logBF*_12_ ≤ 5 is ‘strong’, and *logBF*_12_ *>* 5 is ‘very strong’. It is worth noting that log Bayes factors can be directly converted to posterior probabilities; the posterior probability of model *M*_1_ relative to model *M*_2_, *p*_12_ is given by 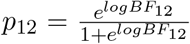 (or 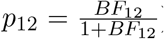), meaning that a *logBF*_12_ = 3 (i.e. between ‘positive’ and ‘strong’ evidence) corresponds to *p*_12_ ≈ 0.95 (Table 1). The key assumption here is that only two models are under consideration and that their prior probabilities are 0.5, such that they cancel out in the Bayes factor ratio.

**Table 1:** Correspondence between log Bayes factors between two competing models or hypotheses (*logBF*_12_) and posterior probabilities (*p*_12_). This equivalence is valid under the assumption that two models are being compared and that their respective prior probabilities are 0.5.

| Degree of evidence in (Kass and Raftery, 1995) | $\log BF_{12}$ | $p_{12}$ |
| --- | --- | --- |
| ‘not worth more than a bare mention’ | 0 | 0.500 |
| ‘positive’ | 1 | 0.731 |
| ‘strong’ | 3 | 0.953 |
| ‘very strong’ | 5 | 0.993 |

Another Bayesian approach for hypothesis testing consists of measuring statistical support for the effect size for a parameter of interest (Kass and Raftery, 1995, Keysers et al., 2020, Morey et al., 2016, van de Schoot et al., 2021). We propose to use this approach for detecting episodic evolution by considering the FLC model, and a well-defined hypothesis; the ratio of evolutionary rates of foreground and background branches is at least 1, meaning an episodic increase in the evolutionary rate. Statistical support is quantified by comparing the posterior and prior odds for the hypothesis of interest, which we refer to here as ‘Bayes factors on effect size’.

Finally, Bayesian model averaging is an approach where a pool of models are considered and estimates of all parameters, including evolutionary rates, are weighted averages of the models, and with the model weights corresponding to their posterior probability. Molecular clock model averaging exists for selecting uncorrelated relaxed clock models (Baele et al., 2013, Li and Drummond, 2011). The random local clock model (RLC) is also a model averaging approach, where nodes can undergo evolutionary rate changes (Drummond and Suchard, 2010). The number and location of evolutionary rate changes is inferred, without the need for their *a priori* definition, as is the case for the FLC. Although it is conceivable that the RLC may be able to identify episodic evolution, its objective is to identify all rate changes, which is fundamentally different to comparing a small subset of models, as in ‘Bayes factors on model comparisons’ or in ‘Bayes factors on effect size’. Moreover, in the RLC evolutionary rates cannot be shared by non-neighbouring branches, meaning that some forms of the FLC (e.g. Fig. 1 C) are not considered.

## 2 Results

### 2.1 Simulating episodic evolution

We conducted a simulation study to demonstrate the performance of Bayesian approaches for detecting episodic evolution. Our simulations utilised 100 phylogenetic trees drawn from the posterior distribution of empirical SARS-CoV-2 data by Tay et al. (2022). These trees contain 179 tips that represent curated full genomes collected between December 2019 and August 2021, and include 20 samples from each of the first four VOCs (Alpha, Beta, Gamma, and Delta) and a global sample of the diversity of the virus at the time. We simulated evolutionary rates along these trees according to the FLC model using NELSI (Ho et al., 2015), where the background branches had an evolutionary rate of 7 *×* 10^−4^ substitutions per site per year, similar to previous estimates for this virus (Boni et al., 2020, Duchene et al., 2020), and the stem branches leading up to VOCs (foreground) had evolutionary rates of 1, 1.5, 2, 3, 5, or 10-fold that of the background (a 1-fold rate increase is equivalent to the SC). For each of the 100 trees, we set each of the six foreground rate increases and simulated a sequence alignment with 29,903 nucleotides, resulting in 600 synthetic alignments.

We fit three molecular clock models to each simulated data set under a Bayesian phylogenetic framework: the SC, the UCGD (uncorrelated relaxed-clock with an underlying gamma distribution; Drummond et al. (2006)), and an FLC configured such that the foreground branches are the stems leading up to VOCs (see Fig. 1c). For the relaxed clock model we chose the UCGD over the more popular UCLD (lognormal distribution), because previous empirical analyses favoured the UCGD for this SARS-CoV-2 data (Tay et al., 2022). We sampled the posterior distribution using Markov chain Monte Carlo (MCMC) as implemented in BEAST 1.10 (Suchard et al., 2018).

We inspected the posterior distribution of the ratio of evolutionary rates of foreground and background branches for the analyses using the FLC model. For the SC simulations, six out of the 100 replicates had a 95% credible interval that excluded a value of 1.0 used to generate the data, which approximates the expected 5% if 95% credible intervals are interpreted as frequentist confidence intervals, and in accordance with the Bernstein-von Mises theorem (see van der Vaart et al. 2017). As expected, increasing the evolutionary rate of foreground branches resulted in fewer replicates where the 95% credible interval included a value of 1.0.

An important trend in these results is that the precision of the estimates improved with increasing foreground evolutionary rates. Indeed, estimates of the foreground to background evolutionary rate ratio are much more uncertain in simulations under the SC (i.e. the simplified case of the FLC), compared to that where the foreground branches had rates were 10-fold higher than the background (SC versus FLCx10 in Fig. 2). A likely cause of this pattern is that increasing the evolutionary rate for foreground branches results in overall more informative sequence data sets.

**Figure 2:**
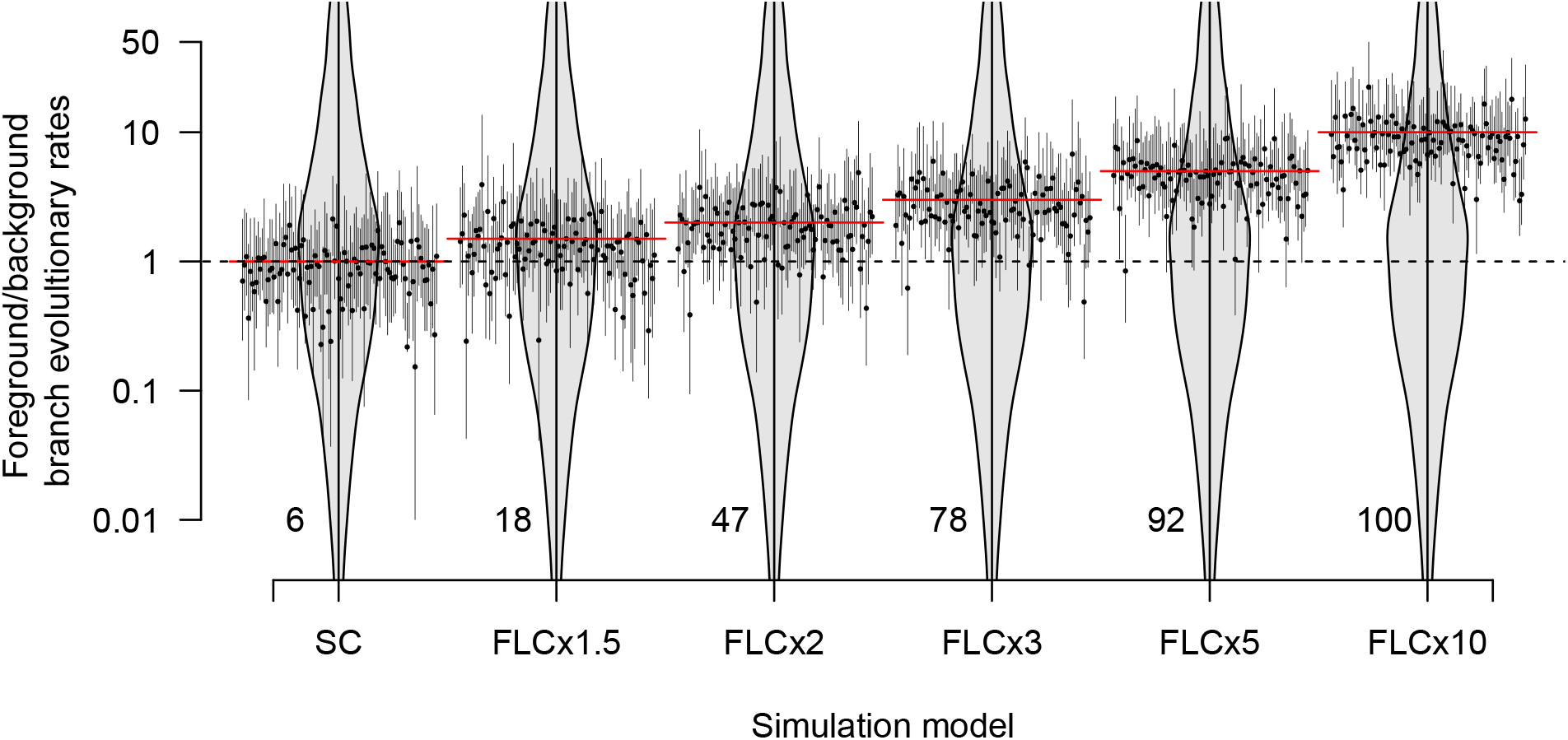
Posterior estimates of the foreground to background evolutionary rate ratio (foreground/background) for all simulation scenarios. SC corresponds to simulations under a strict clock and with foreground and background rates being the same. FLCx1.5 stands for simulations under a fixed local clock, where the foreground branches have an evolutionary rate that is 1.5-fold higher than that of the background, with similar nomenclature through to FLCx10. Each black vertical line denotes the posterior 95% credible interval for each of the 100 simulated data sets and black dots represent the means. The dashed horizontal line represents a foreground/background evolutionary rate ratio of 1.0, such that they are equal. Red lines across each simulation model correspond to the true value used to generate the data. The grey violin plots in the background are used to show the resulting prior on the evolutionary rate ratio. Here we used a Γ(*κ* = 0.5, *θ* = 0.1) prior (where *κ* and *θ* are the ‘shape’ and ‘scale’ parameters and the mean is *κθ*) for the evolutionary rate of both sets of branches, which is relatively uninformative and with the foreground/background ratio centred at 1.0. The numbers at the bottom of each violin are the number of simulation replicates for which the 95% credible interval did not include the foreground/background evolutionary rate ratio of 1.0, which would indicate that no episodic evolution was detected.

The degree of uncertainty in estimates has important consequences for statistical support for episodic evolution. The number of data sets for which the 95% credible interval excluded a value of 1.0, meaning that foreground and background evolutionary rates were different, increased with the degree of episodic evolution. Notably, this was the case for 18 out of 100 simulations under the FLCx1.5, and continues to increase up to 100 for those with FLCx10 (Fig. 2).

### 2.2 Bayesian molecular clock model selection using log Bayes factors on model comparisons

For each simulated data set we estimated the log marginal likelihood using generalised stepping-stone sampling (Baele et al., 2016). We ranked the models and selected them according to the highest log marginal likelihood. An important consideration, however, is the variation in the log marginal likelihood calculations. In particular, selecting a model with a log Bayes factor of 1 (‘positive evidence’) can be unreliable if the uncertainty in log marginal likelihoods is a few log units. To investigate uncertainty in log marginal likelihoods we randomly selected a subset of our simulations and repeated the log marginal likelihood estimations ten times under the three molecular clock models fit to the data (SC, FLC, and UCGD). In Fig. 3(a) we show the resulting repeatability plot, that demonstrates the difference in log marginal likelihoods for two independent runs. We also calculated the average of the absolute difference in log marginal likelihoods between the two randomly selected estimates for each of the three molecular clock models, 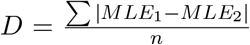, where *MLE*_1_ and *MLE*_2_ are two replicates of log marginal likelihood calculations and *n* is the number of simulated data sets (Baele et al., 2016). The *D* statistic ranged from 0.44 for the FLC to 1.48 for the UCGD. We also calculated the standard error of the log marginal likelihoods as 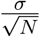, with N=10 replicate log marginal likelihood calculations. In Fig. 3(b) we show the log marginal likelihood for 10 replicates per data set analysed under each of the three clock models. The widths of the 95% confidence intervals (calculated as the mean *±* standard error *×* 1.96) around the log marginal likelihood were on average 1.07. As such, a log Bayes factor of 3 (‘strong’ evidence) seems sufficiently high to distinguish between molecular clock models in light of the uncertainty in these estimates. Note that a log Bayes factor of 3 corresponds to a posterior probability of about 0.953 of the best-fitting model vs. the second best (*posterior* 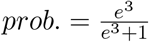).

**Figure 3:**
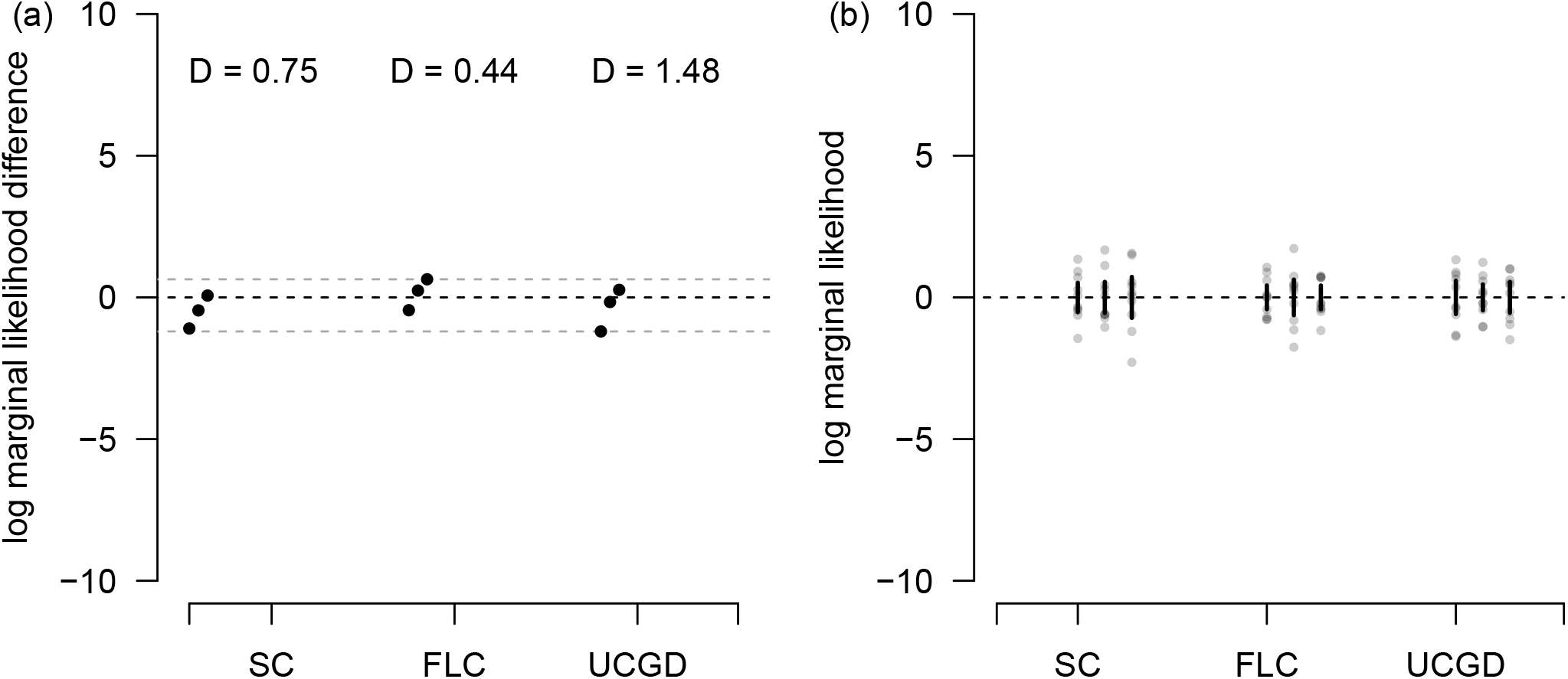
Uncertainty in log marginal likelihood estimates. We repeated the log marginal likelihood estimations for a subset of data sets ten times, analysed under the three molecular clock models, SC, FLC, and UCGD, along the x-axis. In (a) we show the repeatability plot, where each point is the difference in log marginal likelihood between two randomly chosen replicates. The black dashed line corresponds to a value of 0, where there is no difference in log marginal likelihoods and the grey dashed lines are for the maximum and minimum values observed. In each case we show the repeatability statistic, 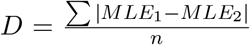, where *MLE*_1_ and *MLE*_2_ are two replicates of the log marginal likelihood estimates and *n* is the number of replicate data sets. In (b) we show the log marginal likelihoods for the ten replicates per data set and per molecular clock model, with the mean value subtracted in each case so that values are centered at 0. Each point is a log marginal likelihood estimate, with the clouds corresponding to replicate estimates for a simulated data set. The solid vertical lines denote the span of the 95% confidence interval, calculated as the mean *±* standard error *×* 1.96. The dashed horizontal line, again denotes a log marginal likelihood value of 0.

In our simulations under the SC we found that no data sets supported the FLC or UCGD. As expected, increasing evolutionary rates along foreground branches resulted in more data sets preferring the FLC. However, a foreground evolutionary rate of at least three-fold that of the background was necessary for most simulations to favour the FLC with ‘strong’ log Bayes factor support (*logBF >*3). For a five-fold evolutionary rate increase, 83 out of 100 simulations selected the FLC, with ‘very strong’ support. In simulations with a ten-fold evolutionary rate increase 97 of 100 simulations displayed at least ‘very strong support’ for the FLC.

### 2.3 Assessing statistical evidence using Bayes factors on effect size

For our analyses under the FLC, we quantified evidence in favour of episodic evolution by comparing the posterior and prior odds, the Bayes factor on effect size. We consider the hypothesis of episodic evolution as the situation where the evolutionary rate of foreground branches (*r*_*f*_) is higher than that of the background branches (*r*_*b*_). Because *r*_*f*_ and *r*_*b*_ are parameters of the model, we treat their ratio (or difference) as the ‘effect size’. The prior probability for the hypothesis is *q*(*r*_*f*_ *> r*_*b*_) and its prior 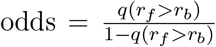. Our prior for both *r*_*f*_ and *r*_*b*_ is a gamma distribution, Γ(*κ* = 0.5, *θ* = 0.1) (where *κ* and *θ* the ‘shape’ and ‘scale’ parameters, respectively, and the mean is *κθ*=0.05), which results in *q*(*r*_*f*_ *> r*_*b*_) = 0.5 and prior odds = 1.0. The posterior probability in favour of episodic evolution is *p*(*r*_*f*_ *> r*_*b*_), with posterior 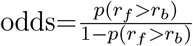. To calculate the Bayes factor we take the posterior and prior odds ratio, which we present in a logarithmic scale to facilitate comparison with other methods. The correspondence between log Bayes factors and posterior probabilities from Table 1 also applies under the condition that the prior probability of the hypothesis in question is 0.5, such that the prior odds are 1.0.

We found that the log Bayes factors on effect size had higher sensitivity than using log Bayes factors on model comparison. For the data simulated under the SC, 37 out of 100 simulations had a positive log Bayes factor (*logBF >* 0, ‘not worth more than a bare mention’), but only 4 displayed ‘strong’ (*logBF >* 3) evidence for episodic evolution. In our simulations for which the foreground branch rates were three-fold higher than that of the background (FLCx3) we found that 82 out of 100 cases had ‘strong’ support for episodic evolution (see Fig. 5 and Table 2). For higher evolutionary rate increases almost all simulations had ‘very strong’ support, at 91 for those with a five-fold rate increase and 99 for those with a ten-fold rate increase.

**Table 2:**
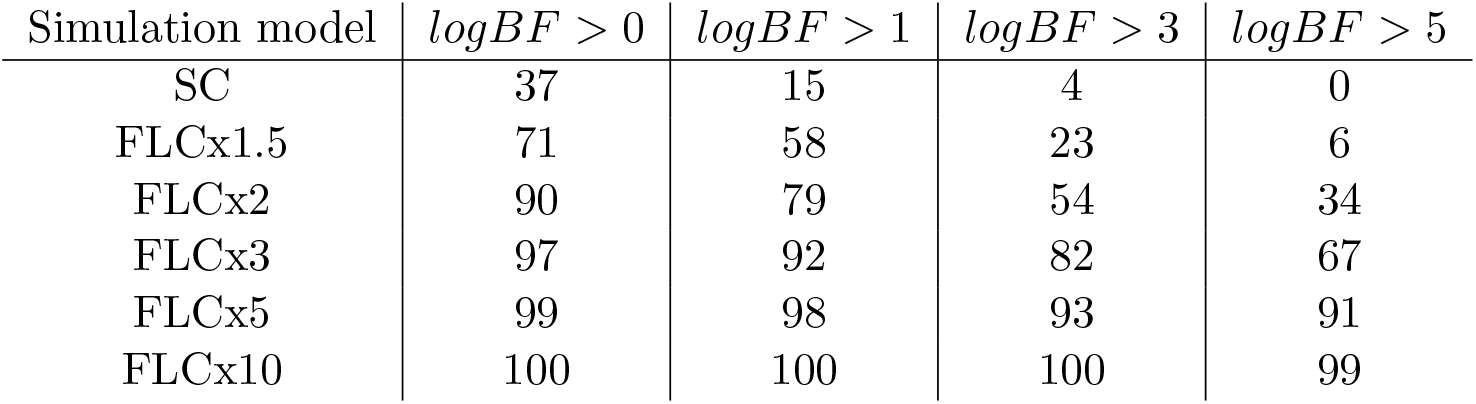
Number of simulated data sets (out of 100) with different degrees of support for episodic evolution according to log Bayes factors on effect size (see also Fig. 5). All data sets were analysed using the FLC molecular clock model, where episodic evolution occurs when the foreground branches have higher evolutionary rates than the background branches. The models used to simulate the data are shown in each row and match those described in Fig. 2.

| Simulation model | $\log BF > 0$ | $\log BF > 1$ | $\log BF > 3$ | $\log BF > 5$ |
| --- | --- | --- | --- | --- |
| SC | 37 | 15 | 4 | 0 |
| FLCx1.5 | 71 | 58 | 23 | 6 |
| FLCx2 | 90 | 79 | 54 | 34 |
| FLCx3 | 97 | 92 | 82 | 67 |
| FLCx5 | 99 | 98 | 93 | 91 |
| FLCx10 | 100 | 100 | 100 | 99 |

**Figure 4:**
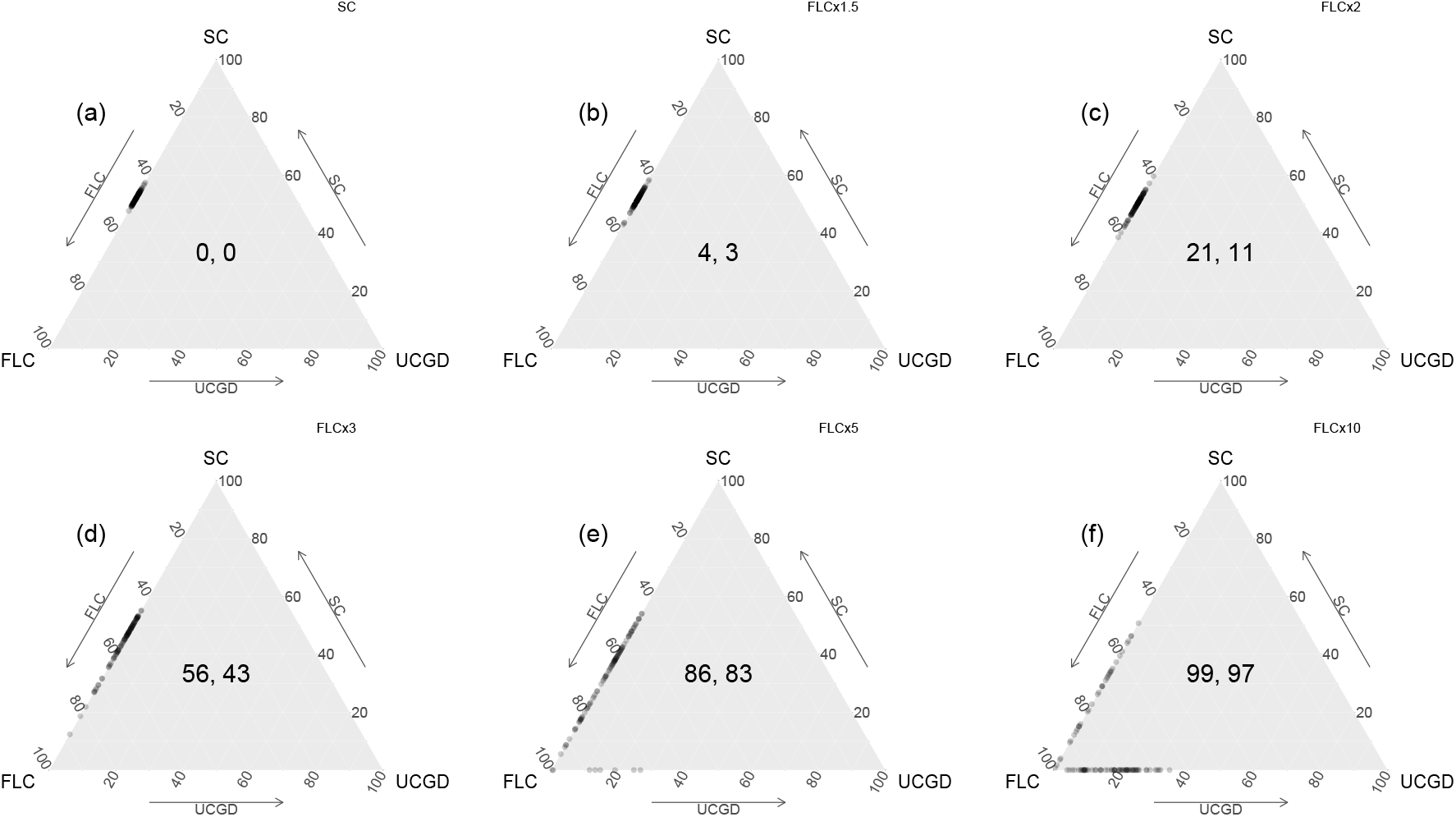
Ternary plots comparing the estimated log marginal likelihoods for the three molecular clock models, SC, FLC, and UCGD. Labels (a) through (f) correspond to molecular clock models used to simulate the data and match those described in Fig. 2. Each axis of the simplex represents the percentage support for a particular model, where each point indicates the level of support for each simulation. A value of 100 shows full support while 0 shows no support for a given model. The numbers in the centre display the total simulations where there is ‘strong’ evidence that the FLC has a higher (log) marginal likelihood than the other models (log Bayes factor *>* 3), and those in which the log Bayes factor *>* 5, implying ‘very strong’ evidence.

**Figure 5:**
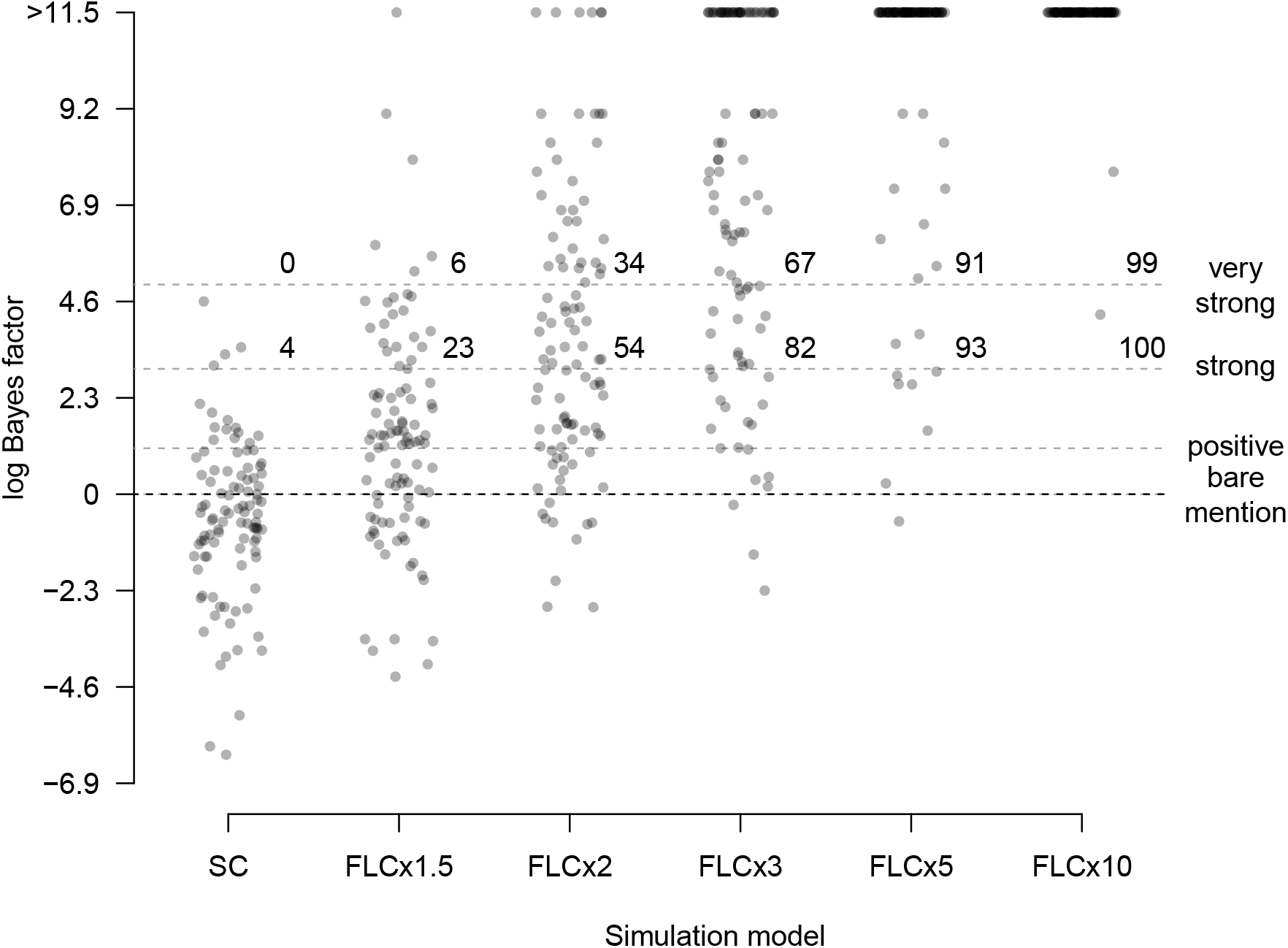
Statistical evidence quantified as log Bayes factors on effect size, here corresponding to the posterior and prior odds of episodic evolution. All data are analysed under the FLC molecular clock model and episodic evolution occurs when the foreground branches have higher evolutionary rates than the background branches. The models used to simulate the data are shown in the x-axis and match those described in Fig. 2. Each point represents the log Bayes factor (y-axis) of a simulated data set (100 per simulation model). The dashed horizontal lines display degrees of support according to log Bayes factors (*>* 0 for ‘not worth more than a bare mention’, *>* 1 ‘positive’, *>* 3 ‘strong’, and *>* 5 ‘very strong’). The numbers above the dashed lines labelled ‘strong’ and ‘very strong’ correspond to the number of points above them, meaning the number of simulations with at least such support.

### 2.4 Prior sensitivity

An important, but often overlooked, aspect of Bayesian phylogenetic analyses is sensitivity to choice of the prior distribution (Gao et al., 2023, Moore et al., 2016). This is particularly relevant for (log) Bayes factors on effect size, but is also likely to impact the (log) marginal likelihood and thus (log) Bayes factor on model comparisons. We randomly selected 25 simulated data sets and repeated the FLC analyses with a different prior on the evolutionary rate as Γ(*κ* = 0.001, *θ* = 1000) for both, foreground and background branches. Under this prior, the mean evolutionary rate is *κθ* = 1.0. Recall that our initial prior for this parameter is Γ(*κ* = 0.5, *θ* = 0.1), with mean *κθ* = 0.05. Both priors are relatively uninformative and do not have high prior probabilities for episodic evolution, at 0.38 for Γ(*κ* = 0.001, *θ* = 1000) and 0.5 for Γ(*κ* = 0.5, *θ* = 0.1). However, the commonly used Γ(*κ* = 0.001, *θ* = 1000) has more weight on low values and its prior odds are not 1.0, as is the case for Γ(*κ* = 0.5, *θ* = 0.1), such that the equivalency between log Bayes factors and posterior probabilities in Table 1 does not hold.

The log Bayes factors on effect size using both prior parameterisations were slightly lower for the Γ(*κ* = 0.5, *θ* = 0.1) prior, with at an average of -0.46 log units difference (Fig. 6(a)). This pattern can be explained by the fact that the prior odds for this prior are 1.0, while those for Γ(*κ* = 0.001, *θ* = 1000) are about 0.61, meaning that the latter can increase the (log) Bayes factor, when the posterior is robust to the prior. The differences between log Bayes factors on model comparisons were larger, with an average of 15.53 log likelihood units, and with the Γ(*κ* = 0.5, *θ* = 0.1) yielding consistently higher values (Fig. 6(b)), which probably occurs because this prior is less diffuse. The finding that the prior on the evolutionary rate has a stronger impact on the log Bayes factor on model comparisons than on effect size is not unexpected, as log Bayes factors on models that differ in their dimensions tend to be sensitive to the prior, even if the posterior itself is not directly affected (Lartillot, 2023).

**Figure 6:**
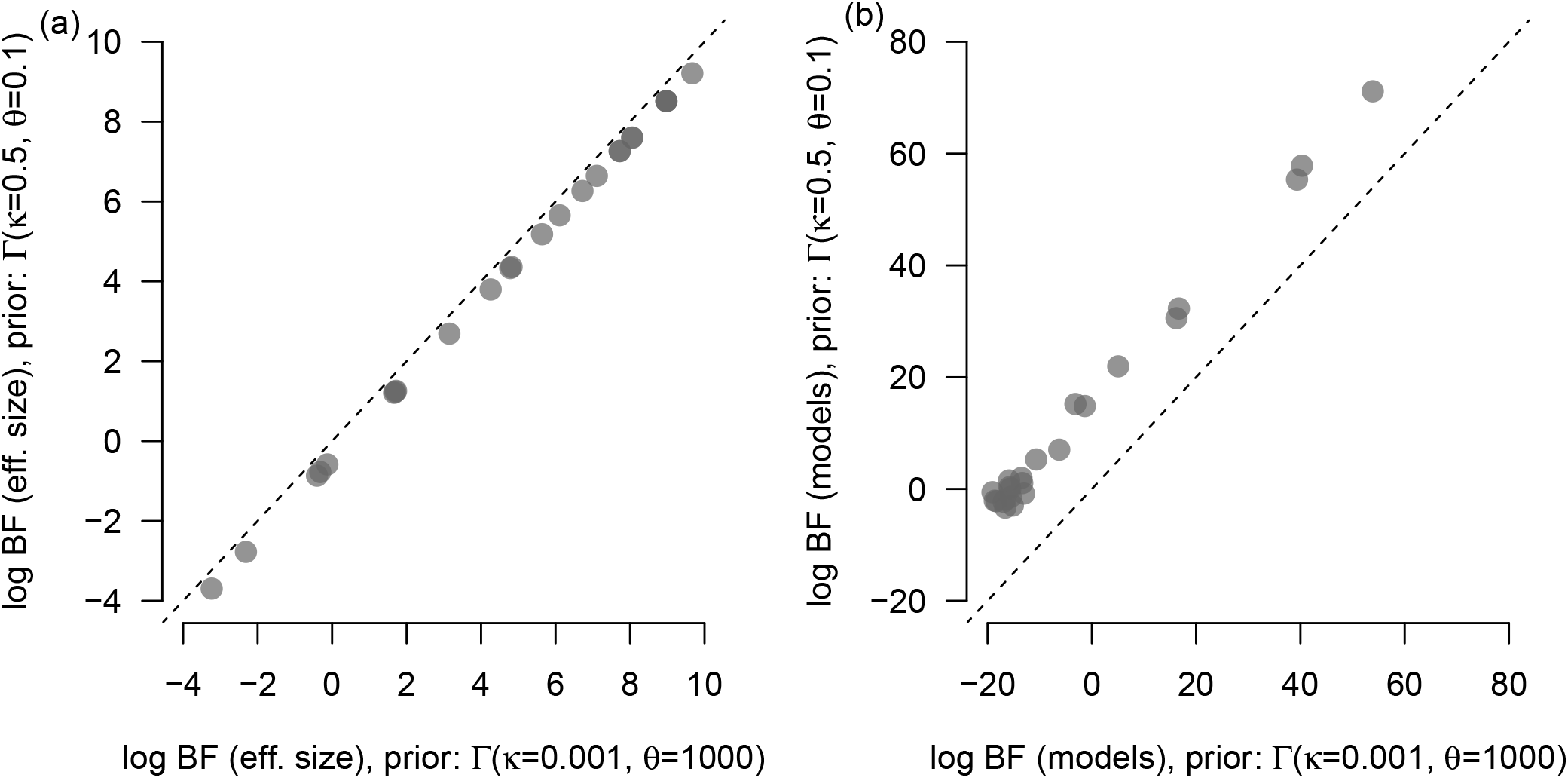
Impact of the prior on evolutionary rate on log Bayes factors (log BF) on effect size and on log Bayes factors for model comparisons (FLC vs. SC). In (a) we show the difference in log Bayes factors for analyses under the using the prior Γ(*κ* = 0.5, *θ* = 0.1) (as used throughout this study) and Γ(*κ* = 0.001, *θ* = 1000) on the evolutionary rate, with *κ* corresponding to the ‘shape’ and *θ* to the ‘scale’ parameters. In (b) we show the corresponding difference in log Bayes factors for model comparisons (models). Each point is for a simulated data set analysed and both prior configurations. The dashed line denotes y=x, where the log Bayes factors for both priors would be identical.

In Fig. 7 we show the prior and posterior for two simulation replicates under the SC and for the FLC where the foreground rate was 3-fold higher than the background (FLCx3). The posterior distributions are nearly identical for either prior, but there is some variation in the log Bayes factors. However, both display a similar degree of support for episodic evolution, with no support for the SC and ‘very strong’ support for the FLCx3.

**Figure 7:**
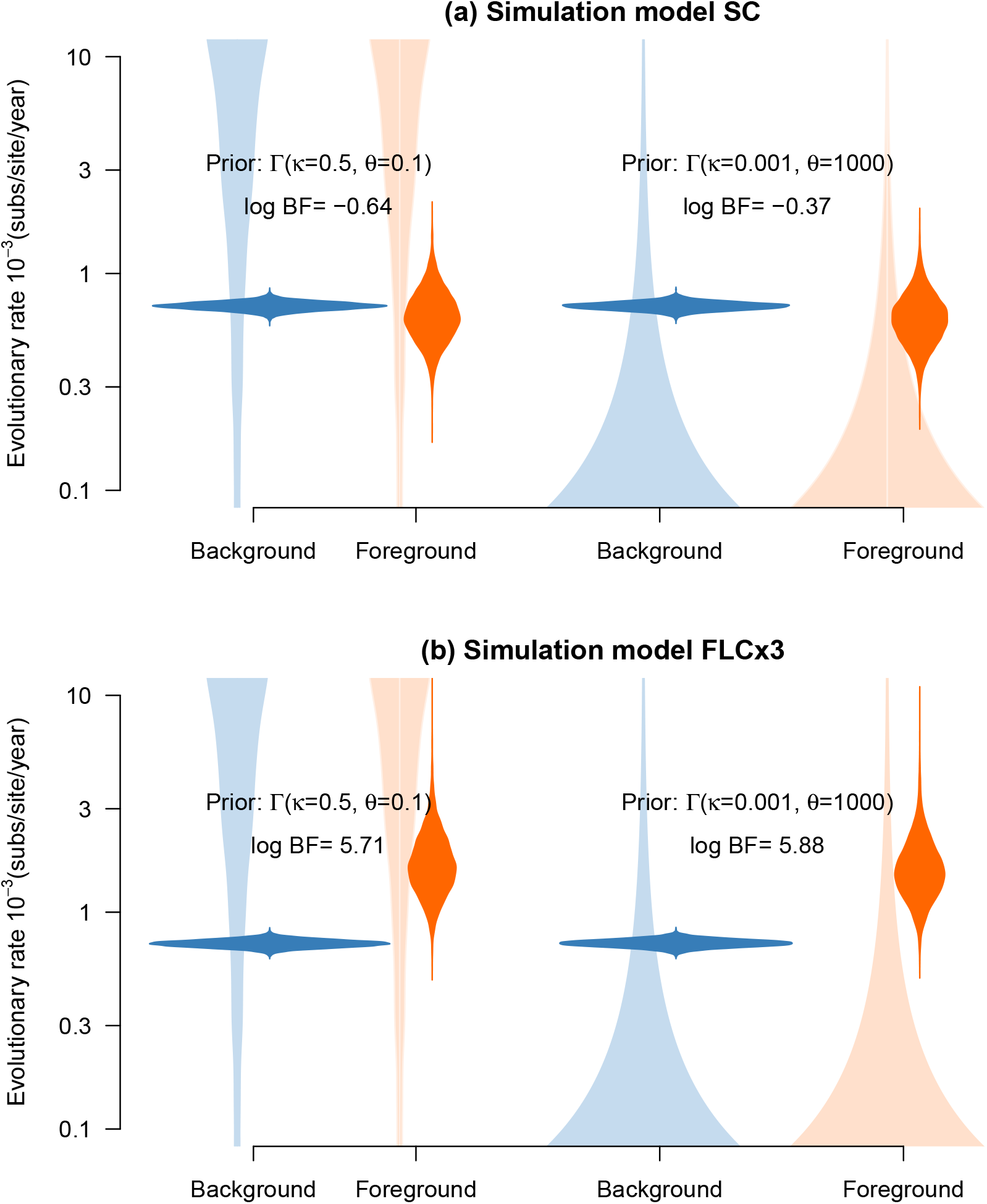
Prior posterior densities for background and foreground branch evolutionary rates under the FLC. The colours match those branches in Fig. 1c. The light blue densities correspond to two possible priors for the background branches, either Γ(*κ* = 0.5, *θ* = 0.1), used in most of our simulations, or Γ(*κ* = 0.001, *θ* = 1000), for assessing prior sensitivity (*κ* and *θ* are the shape and scale parameters). The light orange densities represent the prior on foreground branches, which is identical to the prior on background branches in light blue. The densities in darker shades correspond to the posterior distributions. The log Bayes factor on effect size (log BF) is shown in each case, and is calculated as the posterior and prior odds ratio of an increase in the evolutionary rate (i.e., where foreground branches have higher rates than the background). The top row (a) is for a data set simulated under a strict clock (SC) and thus expected to display no increase in evolutionary rates, and the bottom row (b) is a data set where the foreground branches had an evolutionary rate of 3-fold that of the background (FLCx3).

### 2.5 Bayesian model averaging with random local clocks

We assessed the ability of a random local clock (RLC) model (Drummond and Suchard, 2010) to detect branches undergoing episodic evolution. This task is different to assessing support for a small subset of hypotheses, as in the case of (log) Bayes factors on effect size or model comparisons. Indeed, in this Bayesian model averaging approach, the number and location of evolutionary rate changes is inferred, without the need for their *a priori* definition. A key consideration, however, is that evolutionary rates cannot be shared by non-neighbouring branches, meaning that the true model in our simulations is not part of those considered by the RLC. Our expectation for data simulated under the FLC is that nine clocks should be selected, corresponding to a rate change at each VOC stem and crown nodes (see Fig. 1c).

The posterior mass for our analyses under the RLC model tended to be concentrated on larger numbers of clocks with larger evolutionary rate increases, but the maximum number of clocks was consistently inferred to be below our expectation of nine. Indeed, for our simulations where the foreground branch rate was 10-fold that of the background, the RLC detected on average 3.79 clocks (Fig. 8).

**Figure 8:**
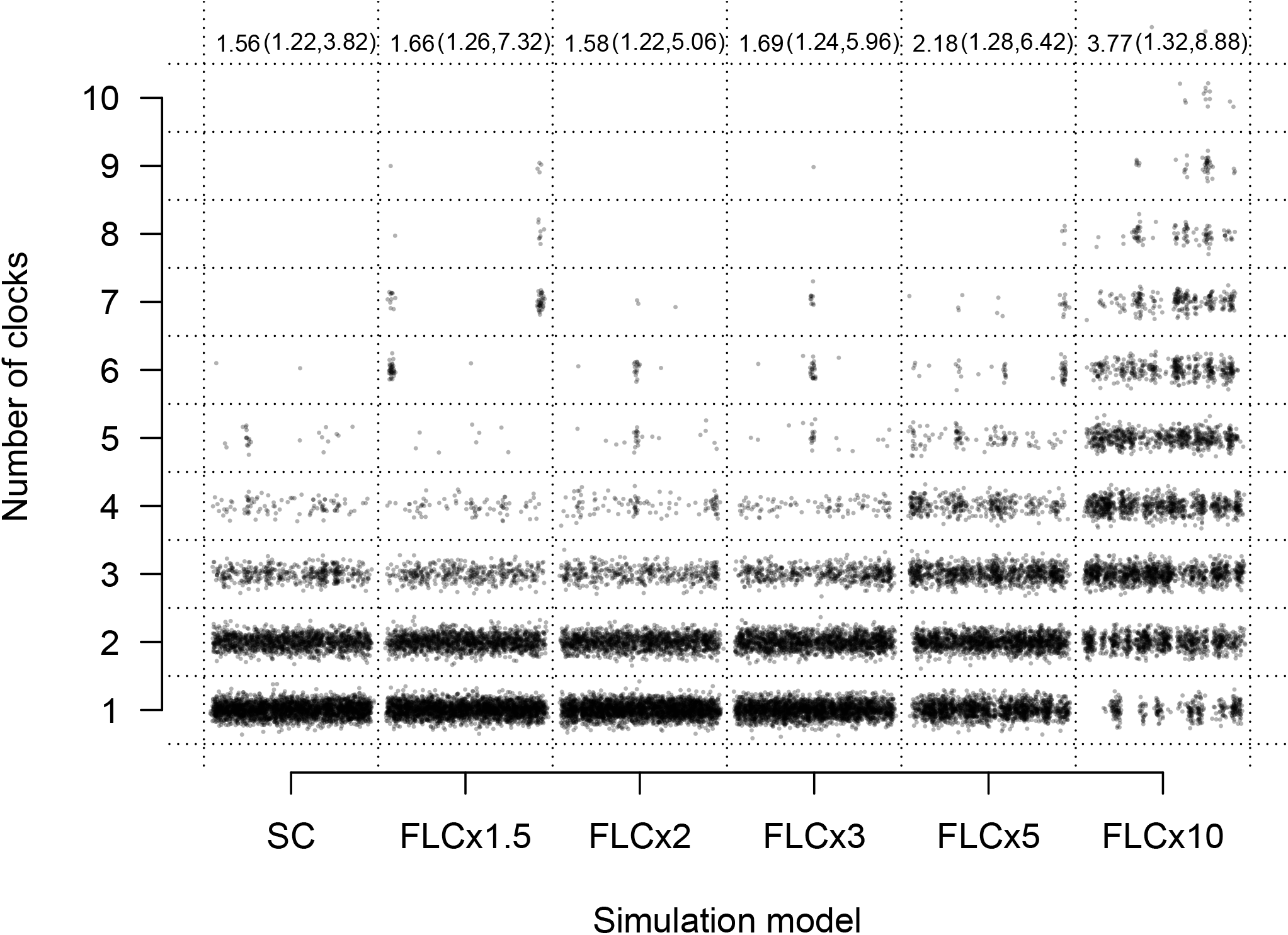
Posterior support for the number of local clocks inferred using Bayesian model averaging under a random local clock (RLC). Each column in the x-axis represents a simulation scenario under the FLC model shown in Fig. 1. The y-axis shows the number of clocks. Each point is a sample drawn from the posterior distribution from the 100 data sets replicate data sets (arranged along each box in the x-axis). Numbers at the top correspond to the mean average number of clocks. Between parentheses we show the highest and lowest mean number of clocks among data sets.

Arguably, a more important question than the number of clocks detected in the RLC is whether any rate changes were detected along the stems or crowns of VOC clades. To investigate this we calculated the posterior probability of rate changes along VOC stem and crown nodes, and took the average over the 100 simulation replicates. We found very low support for rate changes at these nodes (average posterior probability of a rate change ≤ 0.31). It is worth noting, however, that the average posterior probability for a rate change along any of these branches tended to be higher with larger rate changes. For example, the average posterior probability for a rate change in the crown node of Delta was 0.48 for FLCx10, but 4.1 *×* 10^−4^ for FLCx1.5 (Table 3).

**Table 3:** Average posterior probability of evolutionary rate changes along particular branches, using the random local clock model (RLC). Numbers are computed as the posterior probability averaged over 100 simulations in each case.

| Simulation model | Alpha stem | Alpha crown | Beta stem | Beta crown | Gamma stem | Gamma crown | Delta stem | Delta crown |
| --- | --- | --- | --- | --- | --- | --- | --- | --- |
| SC | 0.00 | 0.0043 | 0.00 | 0.0015 | 0.00 | 0.00079 | 0.00 | 0.00060 |
| FLCx1.5 | 0.00 | 0.0076 | 0.00 | 0.011 | 0.00 | 0.00057 | 0.00 | 0.00041 |
| FLCx2 | 0.00 | 0.025 | 0.00 | 0.0057 | 0.00 | 0.0035 | 0.00 | 0.0044 |
| FLCx3 | 0.00 | 0.022 | 0.00 | 0.019 | 0.00 | 0.013 | 0.00 | 0.071 |
| FLCx5 | 0.00 | 0.066 | 0.0060 | 0.081 | 0.00 | 0.11 | 0.00 | 0.22 |
| FLCx10 | 0.050 | 0.30 | 0.060 | 0.24 | 0.02 | 0.31 | 0.02 | 0.48 |

### 2.6 Empirical analyses of SARS-CoV-2 variants of concern

We analysed an empirical SARS-CoV-2 data set to illustrate the application of the methods above. The data set was curated in a previous study by Tay et al. (2022) that displayed ‘very strong’ support for episodic evolution. The data set consists of 20 VOC complete genomes from each of the first four VOCs (Alpha, Beta, Gamma, and Delta) and 99 non-variant genomes, for a total of 179 whole-genome sequences.

We compared the same three models as in our simulations using generalised stepping-stone sampling (note that earlier work used path sampling and stepping-stone sampling to perform log marginal likelihood estimation). We repeated the log marginal likelihood estimation ten times to assess its variation. As expected, the FLC model had the highest support among the clock models tested. If we consider the average log marginal likelihoods over the ten replicates, the log Bayes factor of the FLC with respect to the UCGD (the second best-fitting model) is 6.35 (a posterior probability of *>*0.99), indicating ‘very strong’ support (see Fig. 9). These results were consistent with the log Bayes factor on effect size, at 5.71 (a posterior probability of *>*0.99). The ratio of the evolutionary rate of the foreground branches had a mean of 3.90 (95% credible interval: 2.21 - 8.02). We did not analyse the empirical data under the RLC because such analyses were previously conducted and the RLC did not detect episodic evolution (Tay et al., 2022).

**Figure 9:**
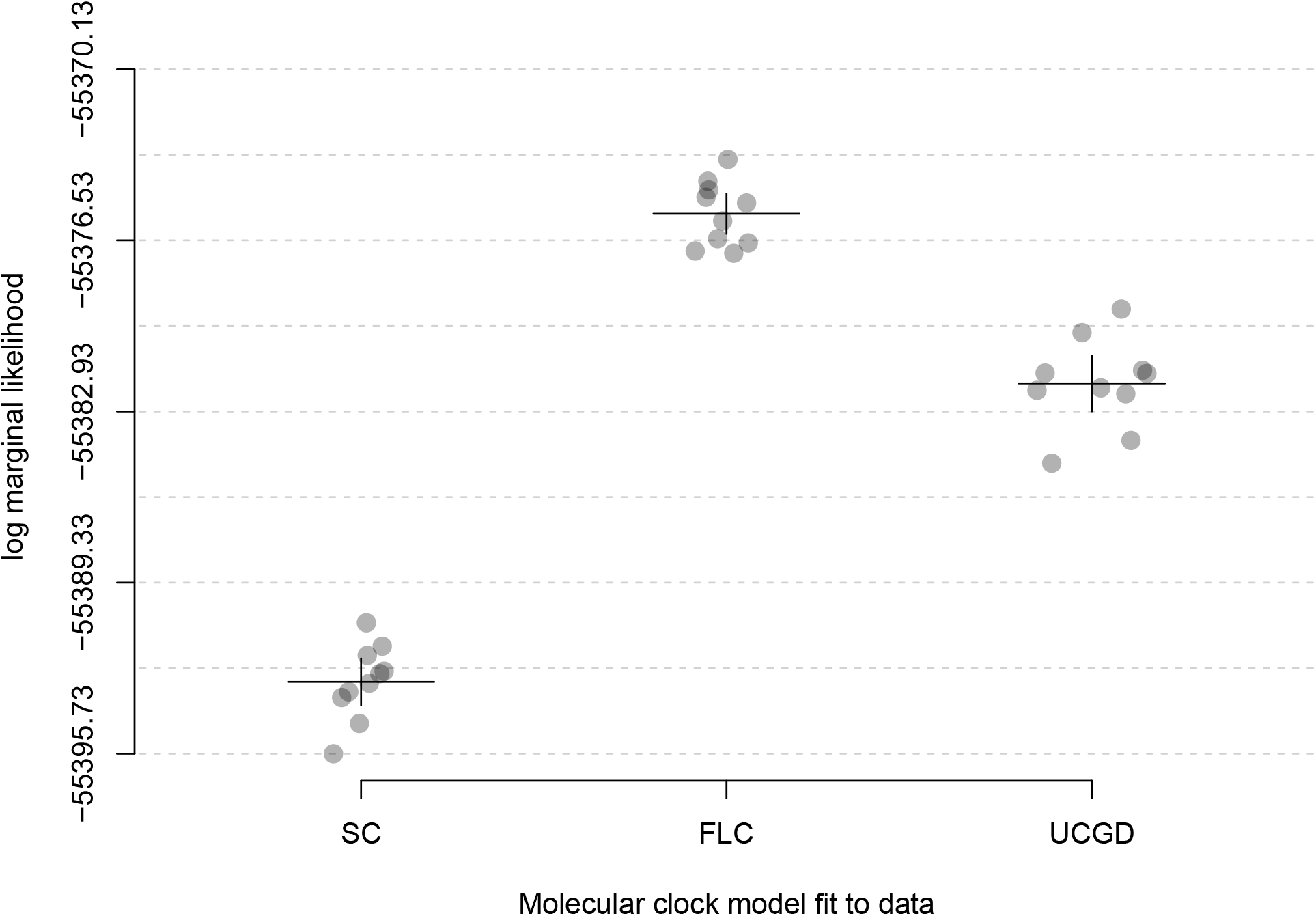
Comparison of the three molecular clocks for an empirical SARS-CoV-2 genome data set, SC, FLC, and UCGD. The x-axis denotes the model used to analyse the data and the y-axis is the log marginal likelihood estimated using generalised stepping-stone sampling. We repeated these calculations ten times to assess their variability. Each point corresponds to a replicate and they have been jittered along the x-axis to facilitate visualisation. The horizontal solid lines represent the average log marginal likelihood across the ten simulations, and the vertical lines are for the 95% confidence interval using the standard standard error. The horizontal dashed lines are spaced by by 3.2 log marginal likelihood units (i.e ‘strong’ evidence.)

We show the inferred SARS-CoV-2 phylogenetic tree under the FLC model in Fig. 10, with branches coloured according to the inferred evolutionary rates. Importantly, some of the branches leading up to VOCs are particularly long, such as that leading up to Gamma, with between 7 and 28 weeks and about 27 expected mutations, consistent with the large number of mutations defining VOCs and their time of emergence (Markov et al., 2023, Martin et al., 2021).

**Figure 10:**
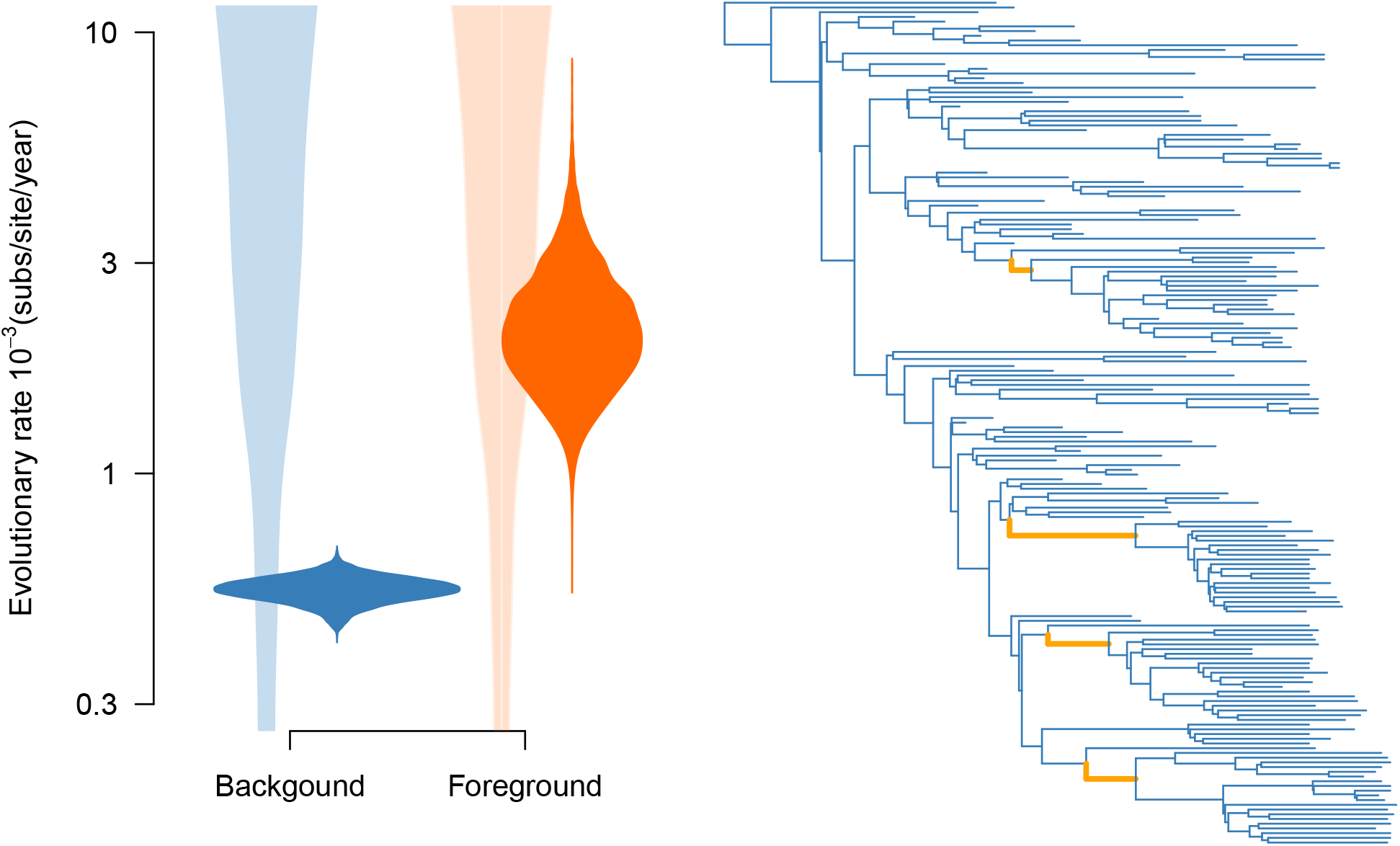
Empirical analysis for SARS-CoV-2 genome data set under the fixed local clock (FLC). The y-axis is the evolutionary rate and the x-axis is for the background and foreground branches, with colours matching those of the tree. The densities in light shades correspond to the prior on the evolutionary rates background and foreground branches, which is a Γ(*κ* = 0.5, *θ* = 0.1) distribution. The densities in darker shades represent the posterior. The log Bayes factor on effect size is 5.71 (‘very strong’ support), and it is calculated as the log ratio of the posterior and prior odds of foreground branch rate being higher than that the background. In the phylogenetic tree, branches coloured in orange denote the stems of the first four variants of concern (VOCs): Alpha, Beta, Gamma and Delta.

## 3 Discussion

Detecting episodic evolution from genomic data can be challenging, particularly in situations where evolutionary rate changes are expected to occur along a small subset of branches, or even a single branch in phylogenetic trees. Our simulations are necessarily limited to one specific scenario, the emergence of SARS-CoV-2 VOCs, but they provide important guidelines for detecting episodic evolution in other organisms and outbreaks. The signal in the data is a key consideration for the sensitivity of the approaches used here. Consider our simulations where the evolutionary rate of foreground branches was three-fold that of the background (FLCx3). The uncertainty in the ratio of evolutionary rates for both sets of branches is sufficiently high that only 78 out of 100 simulations had 95% credible intervals that excluded a value of 1.0 (indicative of no episodic evolution; Fig. 2). Thus, only about a half of these simulations selected the FLC model with ‘strong’ or ‘very strong’ support (at 56 and 43, respectively; Fig. 4). Using log Bayes factors on effect size yielded higher values, at 82 and 67, respectively (Fig. 5).

In contrast, our simulations with a ten-fold increase in the evolutionary rate of foreground branches with respect to the foreground yielded the lowest uncertainty in the ratio of foreground to background evolutionary rates (Fig. 2). None of the 100 simulations included a value of 1.0 for this statistic and 99 had ‘strong’ support in favour of the FLC over other molecular clock models (Fig. 4). Similarly, all of the simulations had ‘strong’ log Bayes factor support for episodic evolution according to the effect size (Fig. 5). The key factor at play here is the fact that our simulations with higher evolutionary rates for foreground branches tended to have increasing numbers of unique site patterns, meaning more information content (average of 1393 unique site patterns for SC, 1437 for FLCx5, and 1510 for FLCx10), and therefore more power to detect episodic evolution with either method. Other considerations for future studies that may impact our ability to detect episodic evolution may be the total number of sequences, the length of the genome or gene in question, and the number of foreground branches.

Our approach of assessing the effect size using posterior and prior odds to calculate the Bayes factor is practical and computationally efficient. However, the definition of the ‘effect size’ should be made explicit. In our study, the effect size is a convolution of two parameters, rather than a single parameter. A more tractable approach may be to quantify the effect size of an actual parameter of the model, where possible. Alternatively, in a situation similar to the one we explored here, one could consider using a hyperprior for the difference or ratio of two parameters. In any case, verifying the prior support for the hypothesis of interest is essential and straight-forward. For complex models, a simple solution is to run the analyses without data to obtain the marginal prior probability and odds of the hypothesis.

Model testing via log marginal likelihood estimation requires performing additional estimation procedures rather than simply sampling from the posterior. We found it to be more conservative than using prior and posterior odds for detecting episodic evolution, but it is important to note that model selection is much more flexible because it allows one to consider a range of sophisticated molecular clock models, including any parameterisation of the FLC. Moreover, to improve sensitivity one can employ a lower support threshold (e.g. ‘positive’ evidence), under the condition that variation in the marginal likelihood is lower than the level of support required.

The Bayes factors on effect size and on model comparisons used here are designed for situations where a set of branches that represent an evolutionary hypothesis are known beforehand. Such hypotheses can be driven by biological knowledge, as is the case for SARS-CoV-2 VOCs. Identifying branches undergoing episodic evolution with no prior knowledge is a more difficult task. For a simple calculation, consider testing for the presence of episodic evolution on individual branches with a log Bayes factor threshold of *>* 3 (‘strong’ evidence, or a posterior probability of *>*0.95). If there are 356 branches, as is the case here, we would expect about 18 false positive tests (0.05*×*356=17.80). Therefore, blindly scanning for a small subset of branches under episodic evolution would be difficult without incurring in a high number of false positives. This situation likely explains why we did not find the RLC to be sufficiently sensitive to detect evolutionary rate changes along VOC stem branches.

It is also important to note that in our study the correct model used to generate the data is not part of those that the RLC can consider. Clearly, other model averaging approaches that can visit the exact model of in question, in this case the FLC, may have higher statistical power. An interesting approach that also explores a range of models, including local clocks, treats branch-specific rates as a Dirichlet process (Heath et al., 2012), implemented in the RevBayes platform (Höhna et al., 2016).

Other methods that warrant further attention in the context of episodic evolution include those where changes in evolutionary rates are modelled as a compound Poisson process (Huelsenbeck et al., 2000). Recent developments in molecular clock models hold great promise for elucidating complex evolutionary patterns for which a hypothesis is difficult to define *a priori*, such as shrinkage-based RLC models (Fisher et al., 2021), and new MCMC proposals for relaxed clock models (Douglas et al., 2021). Fixed local relaxed clocks (Fourment and Darling, 2018), and additive relaxed clocks (Didelot et al., 2021) are interesting alternatives to the models used here, with the former allowing one to employ multiple relaxed clock models along the phylogeny. Our guidelines and simulation framework will be useful to assess new methods and the evidence for episodic evolution in a range of organisms.

## 4 Methods

### 4.1 Simulations

To simulate our sequence data, we sampled 100 trees from the posterior distribution previously obtained (Tay et al., 2022) under a fixed local clock model with shared stems (labeled FLC here). Under this model, the molecular clock is parameterised with two evolutionary rates, i.e. one for the background branches and another for the foreground branches. The foreground branches are the stems leading up to VOCs and the background branches comprise all other branches, including those that form VOC clades. The premise of this model is that the evolutionary rate of the virus accelerated prior to the emergence of VOCs, but with the VOCs themselves evolving at a rate that is statistically indistinguishable from its ancestral and sister lineages (see Fig. 1).

The background branches were always assigned an evolutionary rate of 7 *×* 10^−4^ substitutions per site per year, similar to previous estimates (Boni et al., 2020, Duchene et al., 2020). For the foreground branches, we assigned evolutionary rates that were 1, 1.5, 2, 3, 5, or 10-fold that of the background branches using the R package NELSI (Ho et al., 2015). Note that a rate increase of 1-fold corresponds to a strict molecular clock model (SC). For each evolutionary rate increase in VOC stem branches, we simulated the evolution of a multiple sequence alignment according to a GTR+Γ_4_ substitution model with 29,903 nucleotides to match the genome length of SARS-CoV-2 using Seq-Gen (Rambaut and Grass, 1997).

### 4.2 Phylogenetic analyses and log marginal likelihood calculations

We analysed each of the 600 simulated data sets using BEAST 1.10 (Suchard et al., 2018), under four molecular clock models: the SC, which has a single parameter, i.e. the evolutionary rate; the UCGD that requires two parameters, i.e. the mean and shape of the gamma distribution, but note that the true number of parameters includes the evolutionary rates across all branches; the FLC that has two parameters for the evolutionary rates: one on the background and one on the foreground branches; and the RLC, where the number and location of local clocks is treated as a random variable (Drummond and Suchard, 2010) and the local clock rates are estimated by multiplying a baseline evolutionary rate. Another popular clock model is the relaxed uncorrelated with an underlying lognormal distribution, UCLD, (Drummond et al., 2006). In earlier analyses of SARS-CoV-2 VOCs we found the UCGD had higher support than the UCLD for these data, and thus chose the UCGD here. We assigned a Γ(*κ* = 0.5, *θ* = 0.1) prior distribution for the evolutionary rates in all models (local clock rates in the FLC, the mean for the UCGD, and the baseline rate of the RLC) and an exponential distribution for the shape parameter of the underlying gamma distribution in the UCGD model (Table 4).

**Table 4:**
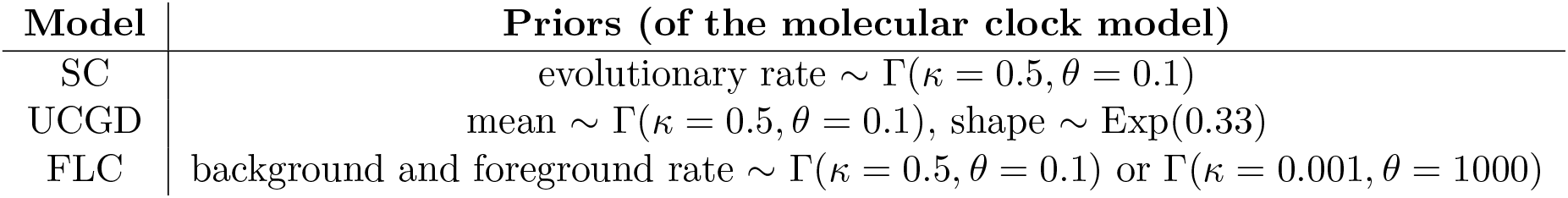
Molecular clock models, parameters and priors used in our Bayesian phylogenetic inference and log marginal likelihood estimation under the three molecular clock models, the SC, UCGD, and the FLC with two evolutionary rates (VOC stem rates as foreground and the rest as background). In the Γ distribution *κ* corresponds to the ‘shape’ parameter and *θ* is the ‘scale’ and the mean value is their product, *κθ*.

| Model | Priors (of the molecular clock model) |
| --- | --- |
| SC | evolutionary rate $\sim \Gamma(\kappa = 0.5, \theta = 0.1)$ |
| UCGD | mean $\sim \Gamma(\kappa = 0.5, \theta = 0.1)$, shape $\sim \text{Exp}(0.33)$ |
| FLC | background and foreground rate $\sim \Gamma(\kappa = 0.5, \theta = 0.1)$ or $\Gamma(\kappa = 0.001, \theta = 1000)$ |

A key aspect of the FLC model here is that foreground branches can become unidentifiable if the topology is co-estimated. To address this problem, we enforced monophyly for each of the four VOCs, with the rest of the tree topology being estimated in the analysis. We also enforced monophyly in the SC, UCGD, and RLC models to ensure that tree space was comparable among the molecular clock models being compared.

For our inference under the different molecular clock models, we specified the GTR+Γ_4_ substitution model and an exponential growth coalescent tree prior to match the simulation conditions. Because we were interested in estimating log marginal likelihoods, we ensured that priors on all parameters were proper probability distributions (i.e. they integrate to one) (Table 5) (Baele et al., 2013). sudo

**Table 5:**
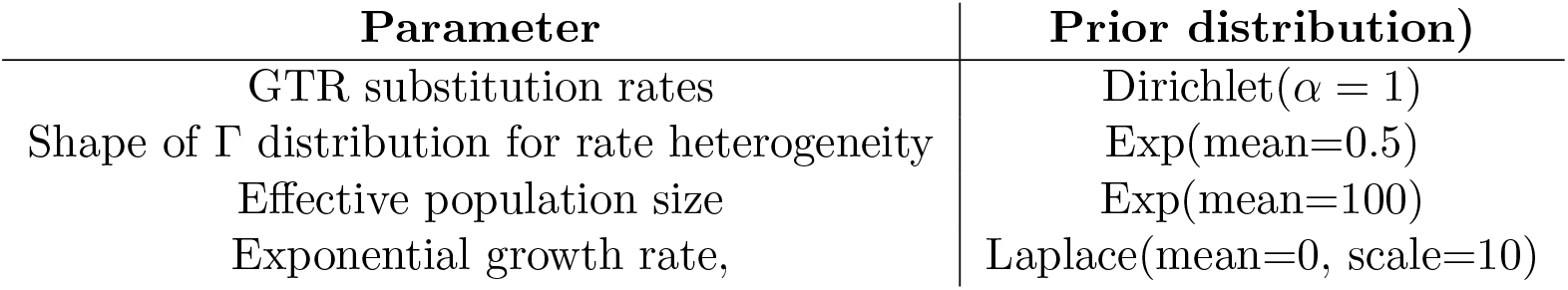
Priors on substitution model and tree prior. The *α* parameter in the Dirichlet distribution is known as the ‘concentration’ and it determines the degree to which the substitution rates in the GTR matrix can differ.

| Parameter | Prior distribution) |
| --- | --- |
| GTR substitution rates | Dirichlet($\alpha = 1$) |
| Shape of $\Gamma$ distribution for rate heterogeneity | Exp(mean=0.5) |
| Effective population size | Exp(mean=100) |
| Exponential growth rate, | Laplace(mean=0, scale=10) |

To sample the posterior distribution, we employed Bayesian inference through MCMC as implemented in BEAST 1.10 (Suchard et al., 2018), running the analyses for a total of 10^7^ iterations, sampling every 10^3^ iterations. We assessed convergence and sufficient sampling from the stationary distribution by verifying that the effective sample size (ESS) was at least 200 for all parameters in the model using Beastiary (Wirth and Duchene, 2022b). In the empirical data analyses we doubled the chain length and reduced sampling frequency accordingly.

We estimated log marginal likelihoods using generalised stepping-stone sampling (Fan et al., 2011). This estimator requires specifying working distributions for all parameters, including the tree (Baele et al., 2016). As working priors for the continuous parameters we used the default in BEAST 1.10 (generally a kernel density estimate of the parameter posterior distribution that is transformed accordingly). For the tree we used the product of exponential distributions determined by intercoalescent times, which is the most general working distribution for the coalescent (Baele et al., 2016).

Importantly, the working distribution requires the same monophyletic constraints on VOC clades, as described above. We specified 100 path steps between the unnormalised posterior and the working distribution, according to equally spaced quantiles from a *β*(0.3, 1.0) distribution. Each path step had a chain length of 10^6^ steps. To assess variation of the log marginal likelihood estimates, we took a subset of simulations and repeated the log marginal likelihood calculations to obtain a repeatability statistic (Baele et al., 2016) and to generate a 95% confidence interval based on the standard error. To verify our log marginal likelihood calculations for this subset of simulations we repeated the calculations with 200 path steps.

### 4.3 Posterior and prior odds and log Bayes factors

To obtain the Bayes factor on effect size, we calculated the posterior and prior odds of episodic evolution. The posterior probability is the proportion of samples from the MCMC for which the evolutionary rate of the foreground branches is higher than that of the background. The posterior odds can then be obtained as explained in the section ‘Assessing statistical evidence using Bayes factors on effect size’. We can use the same approach for the prior probability and prior odds by running the analyses without sequence data. Formally, the probability that one random variable is larger than another is a convolution of probability distributions, with a difficult analytic solution. A more practical approach is to simulate from the corresponding statistical distributions, here Γ, using scientific software, such as R (e.g. Fig. 11).

**Figure 11:**
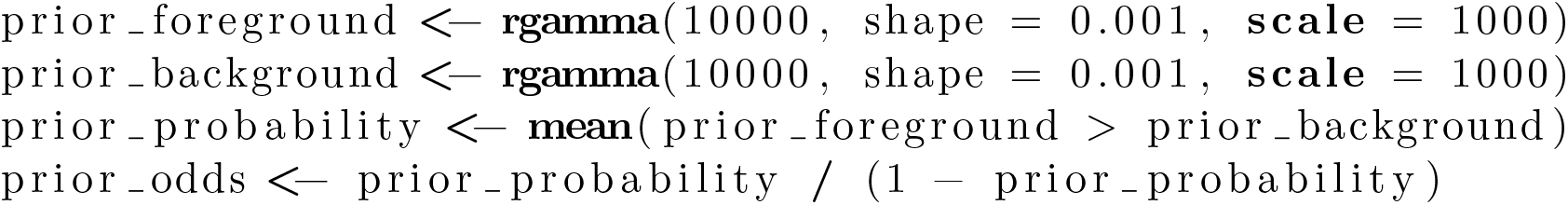
Example of R code to calculate the prior probability and prior odds, where the prior on the evolutionary rates of foreground and background branches is Γ(*κ* = 0.001, *θ* = 1000).

### 4.4 Empirical data of SARS-CoV-2 variants of concern

Our empirical data analyses and simulation settings are based on a data set constructed in a previous study (Tay et al., 2022). These data consist of 99 non-VOC SARS-CoV-2 complete genomes downloaded through Nextstrain (Hadfield et al., 2018) from the GISAID database (Elbe and Buckland-Merrett, 2017, Shu and McCauley, 2017). VOC genomes were included by randomly selecting 20 sequences for the first four reported VOCs (Alpha, Beta, Gamma and Delta), such that the complete alignment comprised 179 whole-genome sequences. To obtain the data, we used the R package GISAIDR (Wirth and Duchene, 2022a). The sequences were aligned using MAFFT v7.49 (Katoh and Standley, 2013) and sites likely to contain sequencing errors were excluded according to those reported previously (De Maio et al., 2020).

## Supporting information

Suppementary Material

## 5 Supplementary Material

Supplementary data are available at Molecular Biology and Evolution online.

## 6 Acknowledgements

The Authors thank two anonymous reviewers and the Editor for helpful comments and suggestions for earlier versions of this manuscript.

JHT and SD were supported by the Australian Research Council (FT220100629) and the Australian National Health and Medical Research Council (grant number 2017284). GB acknowledges support from the Internal Funds KU Leuven under grant agreement C14/18/094 and from the Research Foundation – Flanders (‘Fonds voor Wetenschappelijk Onderzoek – Vlaanderen’, G0E1420N and G098321N). The authors acknowledge efforts by originating and submitting laboratories for the sequence data in GISAID EpiCoV on which our empirical analyses are based. This research was undertaken using the LIEF HPC-GPGPU Facility hosted at the University of Melbourne. This Facility was established with the assistance of LIEF Grant LE170100200.

## 7 Data availability

The data underlying this article are available in GISAID at gisaid.org, and all accession numbers are provided in Supplementary Material online.

## Notes

https://www.dropbox.com/s/s4xwb32yrpviwrk/acknowledgementsTable.tsv?dl=0

### Competing Interest Statement

The authors have declared no competing interest.

### Summary of Updates

This version incorporates changes suggested during peer review, including additional analyses and text.

